# MEG-BIDS: an extension to the Brain Imaging Data Structure for magnetoencephalography

**DOI:** 10.1101/172684

**Authors:** Guiomar Niso, Krzysztof J. Gorgolewski, Elizabeth Bock, Teon L. Brooks, Guillaume Flandin, Alexandre Gramfort, Richard N. Henson, Mainak Jas, Vladimir Litvak, Jeremy Moreau, Robert Oostenveld, Jan-Mathijs Schoffelen, Francois Tadel, Joseph Wexler, Sylvain Baillet

## Abstract

We present a significant extension of the Brain Imaging Data Structure (BIDS) to support the specific aspects of magnetoencephalography (MEG) data. MEG provides direct measurement of brain activity with millisecond temporal resolution and unique source imaging capabilities. So far, BIDS has provided a solution to structure the organization of magnetic resonance imaging (MRI) data, which nature and acquisition parameters are different. Despite the lack of standard data format for MEG, MEG-BIDS is a principled solution to store, organize and share the typically-large data volumes produced. It builds on BIDS for MRI, and therefore readily yields a multimodal data organization by construction. This is particularly valuable for the anatomical and functional registration of MEG source imaging with MRI. With MEG-BIDS and a growing range of software adopting the standard, the MEG community has a solution to minimize curation overheads, reduce data handling errors and optimize usage of computational resources for analytics. The standard also includes well-defined metadata, to facilitate future data harmonization and sharing efforts.

## Introduction

The Brain Imaging Data Structure (BIDS) is an emerging standard for the organization of neuroimaging data^1^. The significance of BIDS is timely: there is increasing availability of open neuroimaging data resources, and strong interest in aggregating large, heterogeneous datasets to harness machine learning techniques and address a new range of scientific questions, with greater statistical power. A single neuroimaging study by itself can represent a large and intricate volume of data, with multiple protocols and modalities, and several categories of participants possibly enrolled in repeated sessions. These aspects are challenging to data organization, harmonization and sharing. The situation is aggravated by the lack of a unique neuroscience standard for digital data across, and sometimes within, modalities such as electrophysiology. Consequently, present data management practices are often based on solutions that do not generalize between labs, or even between persons within the same group. This leads to suboptimal usage of human (time lost retrieving data), infrastructure (data storage space) and financial (limited longevity and value of disorganized data after first publication) resources. Poor or lacking data management strategies also negatively affects the reproducibility of results, even within the lab where the data were collected.

BIDS is a standard to describe the organization of magnetic resonance imaging (MRI) data. It is based on a simple, hierarchical folder structure, with key study parameters documented in text-based metadata files. One benefit is the handling of multiple MRI data sequences, with minimal curation overheads, which reduces the possibility of data-handling errors. An important secondary outcome is the facilitation of interoperability between tools for data analytics, provided that software and pipelines adopt BIDS for data inputs.

We describe here a key extension of BIDS to electrophysiology data. The technical sophistication of MEG makes it the most challenging electrophysiology data type for standardization^2^. For this reason, MEG-BIDS can readily be generalized to electroencephalography, multiunit recordings, and local field potentials. Further to strengthening and rationalizing data management in MEG labs, MEG-BIDS can provide a common structure to present and future large MEG open-data repositories, such as the Human Connectome Project (HCP^3^), the Open MEG Archives (OMEGA^4^), and the Cambridge Centre for Ageing Neuroscience (Cam-CAN^5^). The absence of unique data file format in MEG is compensated by MEG-BIDS’s standard data organization: the sharing and processing of large and complex data hierarchies is simplified, and made compatible and reproducible across tools for data analytics.

To derive the MEG-BIDS specifications, we have combined perspectives from investigators, technical support staff and data managers, with the best-practice guidelines in the field^6^. We also involved the expertise of leading academic software developers for MEG science^7^, including Brainstorm^8^, FieldTrip^9^, MNE^10,11^), and SPM^12^. The proposed MEG-BIDS specifications are presently compatible with these software applications and toolboxes.

## Material & Methods

The MEG-BIDS fields and data organization were defined, bearing in mind the best-practice guidelines for conducting MEG research^6^. The initiative fostered contributions from multiple MEG experts, with a face-to-face discussion at BIOMAG2016, the International Conference on Biomagnetism, where the first incarnation of MEG-BIDS was introduced^13^. The first set of feedback comments and the minutes from further group discussions are publically available (https://groups.google.com/d/msg/bids-discussion/xTHBsGhu0hk/MN25xbxRBwAJ).

A first version of the present manuscript was shared via the preprint server bioRxiv, from where more comments stemming from the community were collected and considered for improvement of MEG-BIDS. A poll survey to probe the interest of the concerned community was also conducted (LINK TO RESULTS).

MEG-BIDS builds on the BIDS hierarchical data structure. For instance data descriptors such as *subject*, *session, technique, run* are BIDS notions that were re-used in MEG-BIDS. Similarly, the simple although extensively used human and machine readable file formats (JavaScript Object Notation [JSON], and Tab Separated Value [TSV] text files) that contributed to the versatility and practicality of documenting metadata elements in BIDS, were expanded with MEG-BIDS. MEG-BIDS employs a straightforward terminology, cautiously defined in line with the general BIDS specifications, although adapted to the unique requirements of MEG. For further reference, the MEG-BIDS specifications are detailed in an open-access online document: https://docs.google.com/document/d/1FWex_kSPWVh_f4rKgd5rxJmxlboAPtQlmBc1gyZlRZM)

The terms used refer to notions that were defined by reaching a consensus amongst the MEG-BIDS contributors and the MEG community. For example, ‘*Subject*’ refers to the scanned participant. Note that from a technical standpoint, MEG is not a *scanning* technique. Yet, we used this terminology for convenience, affinity with other neuroimaging modalities, and to reflect the language used in most MEG labs. A ‘*Session*’ defines a non-intermittent period of time during which the subject is in the scanner. A ‘*Run*’ is a period of time during which empty-room (for noise characterization) or brain activity is recorded continuously, with no interruptions. It is typical with MEG that a *session* consists of multiple *runs*: task instructions can change and/or participants can take a break between runs. The notion of ‘*Task*’ refers to the instructions (and corresponding stimulus material) that are performed by the participant. ‘*Responses*’ is a feature to indicate the recorded behaviour of the subject in relation to the task.

As with the general BIDS specifications^1^, MEG-BIDS file names are constituted by a series of key-value pairs, with multiple possible file types. Some typological aspects are mandatory, while others remain optional, although required to abide to the BIDS guidelines. MEG-BIDS can therefore register data of any kind, including but not limited to task-based, resting-state, and empty-room MEG recordings (e.g., for noise estimation purposes). We emphasize that all of the above notions apply also to EEG, and all modalities of electrophysiology, for which MEG-BIDS serves as template for standardization.

There is no common, open or standard file format in MEG, equivalent to DICOM or NIfTI in MRI. MEG systems manufacturers (CTF, Elekta/Neuromag, BTi/4D Neuroimaging, KIT/Yokogawa/Ricoh, Tristan Technologies, ITAB, KRISS/Compumedics Neuroscan, York Instruments) all cater a vendor-specific format. With MEG-BIDS, unprocessed (raw) data is stored in the native file format (see Discussion for further consideration of common MEG format initiatives) and users can still rely on their preferred data analysis application, possibly provided by the MEG vendor, to browse and read in the MEG file contents. Software can also extract meta information elements from the raw data files. They concern e.g., data collection parameters and other study descriptors, which are eventually transcribed into sidecar JSON files by said MEG-BIDS compatible applications. One major benefit of metadata extraction is the facilitation of subsequent data searches and indexation, without the handling and repeated parsing of large raw data files. Additional relevant files can be included alongside the MEG raw data: some propositions are detailed in the online specifications.

For a given investigator or research group, MEG-BIDS describes a hierarchical structure that descends from a *‘Study’* folder. Multiple *‘Subject’* subfolders contain the data from the participants enrolled. They are arranged by *‘Session’*, each session subfolder containing *‘Run’* folders and eventually, data and metadata files.

The *‘Run’* folder includes a variety of files: MEG recording files in native format, a sidecar JSON document (*_meg.json), a channel description table (*_channel.tsv), and other general BIDS files, such as task events tables (*_events.tsv) that are likely to be specific of each run. *‘Session’* specific files include the coordinates of anatomical landmarks and head-localization coils stored in a JSON document (*_fid.json), optional photographs of the anatomical landmarks and/or head localization coils (*_photo.jpg), fiducials information (*_fidinfo.txt), 3-D scalp digitalization files (*_headshape.<manufacturer_specific_format>) and acquisition times (scans.tsv). The ‘*Subject’* and *‘Study’* specific files are inherited directly from the general BIDS specifications (e.g., participants.tsv). Note that in case of conflict between fields of different runs/sessions, it the inheritance principle should be applied: the description file closer to the data prevails (see Section ‘3.5 The Inheritance Principle’ of the BIDS specifications^1^.

One issue that required special attention was the multiplicity of coordinate systems and units between MEG systems. To impose a unique coordinate system for BIDS based on the subjects’ brain anatomy (e.g., MNI coordinates or equivalent) was an appealing solution, which however would lack generalizability in the MEG practice. MEG data can be collected without anatomical information, such as empty-room noise recordings, which are important to optimal source modeling^6^. MEG-BIDS therefore associates all recordings with a coordinate file defined according to the MEG system used. Again, MEG-BIDS compatible software can read and interpret this information properly.

Akin to MRI, we anticipate that the systematic data organization enabled by MEG-BIDS will be supported by an increasing number of neuroimaging tools, and that more shared data repositories will be organized accordingly. The straightforward design of MEG-BIDS makes it an interoperable common exchange format for transferring data between investigators and community repositories e.g., OMEGA^4^ and OpenfMRI^14,15^. It also facilitates multimodal integration (between MRI, fMRI, MEG, etc), as the data from multiple modalities follow the same organization scheme.

## Open MEG-BIDS datasets

We provide four different publically-available datasets in MEG-BIDS format (~200GB). They are freely available for download from the International Neuroinformatics Coordinating Facility (INCF)’s GitHub: https://github.com/INCF/BIDS-examples/tree/bep008_meg

The MEG-BIDS sample data release includes:

### OMEGA Resting-State samples

Five minutes of eyes-open, resting-state MEG data is available for 5 subjects from The Open MEG Archive (OMEGA)^4^. The data are available from the Brainstorm Tutorial: tutorial MEG resting state & OMEGA database. The first release of data in MEG-BIDS format (~10.5GB) available here: https://box.bic.mni.mcgill.ca/s/omega?path=%2FContributions%20(in%20BIDS%20format)%2Fsample_BIDS_omega

### Brainstorm Auditory Example dataset

Brainstorm Auditory tutorial dataset^8^ (~2.3GB): https://box.bic.mni.mcgill.ca/s/omega?path=%2FContributions%20(in%20BIDS%20format)%2Fsample_BIDS_auditory (released in Public Domain; includes defaced anatomical T1 of participant).

### MNE Sample data

Sample data with visual and auditory stimuli described in^11^: https://drive.google.com/drive/folders/0B_sb8NJ9KsLUQ3BMS0dxZW5nSHM?usp=sharing (released in Public Domain; includes anatomical T1 of participant as well as flash MRI sequences).

### OpenfMRI study ds000117

A multi-subject, multi-modal human neuroimaging dataset of 19 subjects participating in a visual task^16^ (~178GB): https://openfmri.org/dataset/ds000117/. This dataset is used in one of the SPM tutorials, for training purposes: http://www.fil.ion.ucl.ac.uk/spm/doc/manual.pdf#Chap:data:multimodal

## Software

Widely used MEG software packages have already added functionality to support MEG-BIDS:

### Brainstorm^8^

Brainstorm is an application with rich graphical-user interactions and analytic pipeline designs for MEG, EEG, NIRS, and electrophysiology recordings. BIDS-formatted MEG/EEG datasets can be imported automatically into the Brainstorm database, as described in the OMEGA tutorial: http://neuroimage.usc.edu/brainstorm/Tutorials/RestingOmega

### FieldTrip^9^

FieldTrip is an open-source MATLAB toolbox for the analysis of MEG, EEG, and other electrophysiological data. Like most other tools listed herewith, FieldTrip can implement full analysis pipelines, starting from coregistration, preprocessing, time-and spectral analysis, source reconstruction, connectivity and statistics. Among others, FieldTrip has been used for the MEG part of the Human Connectome Project. Since a FieldTrip analysis pipeline is represented as a MATLAB script, its application on BIDS structured data implies that the BIDS details are represented in the analysis scripts that users write.

### MNE^10,11^

MNE (http://martinos.org/mne), whose name stems from its capability to compute cortically-constrained minimum-norm current estimates from M/EEG data. It is a software package that provides comprehensive analysis tools and workflows including preprocessing (Maxwell filtering, ICA, signal space projectors), source estimation (eg. using MNE, beamformers or mixed-norm sparse solvers), time-frequency analysis, statistical analysis including multivariate decoding, and several methods to estimate functional connectivity between distributed brain regions. The core of MNE is written in Python and is distributed under the permissive BSD Licence. MNE will use the BIDS data structure to distribute all its tutorial datasets and the documented analysis scripts. MNE provides Python code to read and write files in BIDS compatible format, as well as summary reports automatically generated via the MNE report command. A preliminary version is available at https://github.com/mne-tools/mne-bids.

### SPM^12^

SPM (http://www.fil.ion.ucl.ac.uk/spm) is a free and open source software written in MATLAB where many widely used methods for the analysis of PET, fMRI and for computational neuroanatomy have been originally developed and implemented. More recently SPM has been extended to perform M/EEG analyses, including topological inference for neurophysiological data, empirical Bayesian framework for source reconstruction and Dynamic Causal Modeling (DCM), an approach combining neural modeling with data analysis. SPM12 includes a library, spm_BIDS.m, to parse and query BIDS-formatted datasets, as well as low-level functions to read/write JSON and TSV metadata files. A complete pipeline for the analysis of a group MEG dataset in BIDS format is presented in preparation.

## Other MEG-BIDS tools

Another set of tools have been developed to generate sidecar MEG-BIDS JSON files, and assist researchers in their evaluation and adoption of the standard. These Python scripts are publically available: https://github.com/INCF/pybids.

For a more detailed description of the MEG-BIDS specification, example datasets, resources and feedback, please visit http://bids.neuroimaging.io.

## Discussion

Although MEG-BIDS is a proposal to establish a standard framework for the organization of electrophysiology data, it does not impose the standardization of the data file format per se. Some initiatives are aiming towards the definition of a new common binary file format for electrophysiology (MNE python group). Akin to DICOM in MRI, one single file format would beneficial, when considering the diversity of native raw file formats in electrophysiology. Yet, MEG data volumes are typically very large (several GBs), hence their duplication into a standard format may be unpractical. Our position is rather to promote MEG-BIDS and ascertain that tools for data analytics continue to be equipped with the necessary readers for all existing vendor formats. We believe the capacity of MEG-BIDS to organise data without requiring a common data format is actually a strength: the standard is flexible in the sense that any data parameters can be extracted and stored as metadata in sidecar json files at the time of creating a new data entry, regardless of the file format for the raw data. Therefore, any new data format for electrophysiology, including emerging standards, is by construction compatible with MEG-BIDS.

Along the same lines, special attention was given to the handling of the various coordinate systems used by the different MEG vendors and toolboxes. MEG-BIDS is also flexible in that respect, as long as the conventions used are characterized and documented in the *_fid.json file. The coordinate systems presently handled by MEG-BIDS are detailed in the Specification document. The coordinate systems used for MEG and EEG sensors, MRI volumes, locations of fiducials, anatomical landmarks and digitized head points, need to be described following this principle, as some are likely to be different.

We aim at extending MEG-BIDS further towards the handling of processed data, which for now and akin to MRI-BIDS, are simply stored in a data Derivatives folder.

MEG-BIDS represents a significant effort towards a common standard for MEG. We anticipate that the MEG-BIDS software ecosystem and the variety of publicly available MEG-BIDS datasets will grow and incentivize the research community towards adoption.

**Figure 1:**
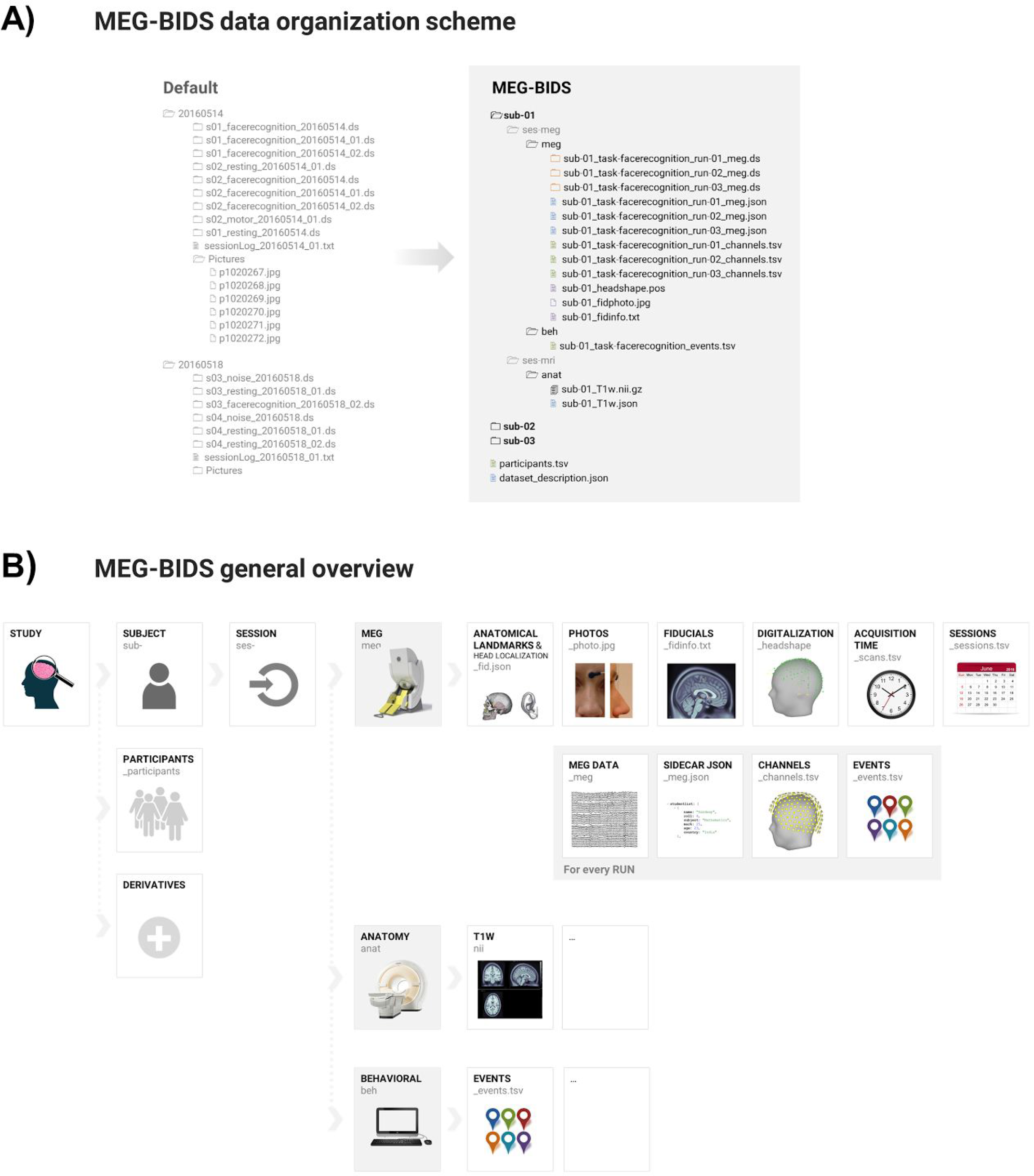
A) MEG-BIDS data organization scheme: Left: a typical default data organization scheme where folder areorganized by date of session and contain different runs for a given participant in a study. *Right*: MEG-BIDS organizes data per study, then participant (subject), followed by modality, then sessions and eventually, runs. Note the sidecar files that are present at all level the data hierarchy, and document conveniently the metadata contents. **B) MEG-BIDS general overview**

## Acknowledgements

This work was supported by the ERC Starting Grant SLAB ERC-YStG-676943 to Alexandre Gramfort, by the EU-H2020 Marie Curie “ChildBrain” Innovative Training Network grant no. 641652 to Robert Oostenveld, by an NWO-VIDI Grant (864.14.011) to Jan-Mathijs Schoffelen, and by a Discovery Grant from the National Science and Engineering Research Council of Canada, the NIH (2R01EB009048-05) and a Platform Support Grant from the Brain Canada Foundation to Sylvain Baillet. The UK MEG community is supported by Medical Research Council grant MR/K005464/1 and The Wellcome Centre for Human Neuroimaging is supported by core funding from the Wellcome [203147/Z/16/Z]. Guiomar Niso received financial support from the AXA Research Fund.

